# Identifying Agronomic, Nutrition and Leaf Vein Quantitative Trait Loci in the Orphan Crop *Gynandropsis gynandra*

**DOI:** 10.1101/2024.09.28.615607

**Authors:** Conor J. C. Simpson, Dêêdi E. O. Sogbohossou, Gregory Reeves, M. Eric Schranz, Pallavi Singh, Julian M. Hibberd

## Abstract

A sustainable food supply is vital for addressing the challenges of a growing global population and climate change. *Gynandropsis gynandra*, a nutritious C_4_ orphan crop native to Africa and Asia, thrives in low-input agricultural systems, making it a valuable candidate for future food security. This crop also serves as a model for studying C_4_ photosynthesis. However, research on its genetic improvement is limited. In this study, two mapping populations from diverse parental lines were used to identify Quantitative Trait Loci (QTL) linked to agronomically relevant traits like plant height, leaf area, flowering time, nutritional content, and photosynthesis. Fifteen QTL were identified, with two affecting both leaf size and flowering time across populations, which can be applied for marker assisted selection for crop improvement. Additionally, QTL linked to C_4_ photosynthesis provide insights into the genetic mechanisms behind this pathway. Overall, this research enhances the potential of *G. gynandra* as a climate-resilient crop.

**One sentence summary:** Leveraging natural variation in *Gynandropsis gynandra* to identify QTL associated with important traits.

## Introduction

At least 250,000 plant species have been identified as edible. Of these, 7000 have been consumed by humans but in 1995 only twelve crops and five animals made up 75% of the world’s food. Moreover, at this time 60% of global calories were provided by just three cereals - rice (*Oryza sativa*), maize (*Zea mays*) and wheat (*Triticum aestivum*)^1^. This bottleneck in agricultural diversity led to interest in underutilized, neglected, or so-called orphan crops^2^ however, there is debate about the use of such terms^3^ and now more inclusive and positive terms such as “opportunity crops” are increasingly being used^4^. Greater use of such species could reduce the demand to adjust conditions to suit to the current limited portfolio of major crops and instead allow crops better adapted to local conditions to be deployed^5^. Thus, increased use of locally adapted species could increase resilience of supply chains and so contribute to improving the lives of the ∼10% of the global population who are undernourished^6^. Better adoption of these species could also help agriculture cope with increasingly unpredictable extreme weather and greater climatic uncertainty that are applying mounting pressure on the global food system^7,8^. Neglected and opportunity-utilized crops are defined as those that have been locally cultivated from sites at which the crop originates, or have become naturalized but been neglected by the global agricultural community^9,10^. Orphan crops are often defined as being rich in nutrient content, are climate resilient, locally available, and economically viable^11–13^. Orphan crops have already been adopted, with examples including quinoa (*Chenopodium quinoa*)^14^ and India’s targeting breeding programme of pearl millet (*Pennisetum glaucum*)^15^.

*Gynandropsis gynandra* of the Cleomaceae family is a relative of Arabidopsis and Brassica crops in its sister family Brassicaceae. It has many common names including Shona Cabbage, Spider Plant, and Cat’s Whiskers. It is grown as an indigenous vegetable crop in parts of Sub-Saharan Africa and Asia and matches the four criteria outlined above^16^. It has recently been recognized as being of high relevance to achieve food security in Africa due to its nutritional content and modelled ability to withstand climate change in Sub-Saharan Africa^17^. *Gynandropsis gynandra* has been identified as a strong candidate for improvement due to its high vitamin C (ascorbic acid) and β-carotene content along with high levels of phosphorus, potassium, calcium, iron, zinc, phenols and flavonoids compared with commercial cultivars of *Brassica oleracea* (var. *capitata* cv. Drumhead) and *Beta vulgaris* (L. cv. Fordhook Giant)^18^. Its seeds are high in polyunsaturated fatty acids and are cultivated in Asia for oil^19^. The active ingredient of PIXALIA (Laboratoires Expanscience) is extracted from *G. gynandra* leaves for acne treatment, and *G. gynandra* has been identified as being of medicinal interest with leaves high in flavonoids such as quercetin and kaempferol^20^ that are thought to have antithrombogenic properties^21^. *Gynandropsis gynandra* extracts have also been reported to contain moderate antifungal and antibacterial activities^22^ and high glucosinolate content, which provides resistance to pests^20^. Lastly, its seeds are well suited for animal feed^19^.

In addition to its practical applications, *G. gynandra* is a model organism for the study of C_4_ photosynthesis. As such it has been used extensively as a system through which genetic determinants of C_4_ gene expression have been investigated^23–27^. Furthermore, recently a full genome sequence for *G. gynandra* was published, revealing their utility in studying genome evolution^28^.

*Gynandropsis gynandra* is grown across a wide geographic range that includes all inhabited continents^19^, and these divergent environments and populations have resulted in a diverse gene pool. For example, resequencing data for fifty-three accessions identified three genetically distinct groups derived from West Africa, Asia and East/Southern Africa, respectively^29^. These accessions exhibited natural variation in multiple economically important traits, including plant height, flowering time, and vitamin content^30^. Local cultivation is a major part of *G. gynandra*’s natural history, as it has been traditionally used by farmers who have independently selected for desirable traits such as stem and leaf colour, plant height, and flowering time^31^. Geographically diverse lines of *G. gynandra* can hybridize, but it is also self-compatible^16^. There is now a rich germplasm collection, with 437 accessions stocked by the World Vegetable Center (https://avrdc.org/). These factors make *G. gynandra* an excellent candidate for pre-breeding methods, such as identifying Quantitative Trait Loci (QTL) for use in Marker Assisted Selection (MAS).

Crossing highly divergent lines enables segregating mapping populations from which QTL can be identified, providing insight into trait inheritance and targets for marker assisted selection to enhance germplasm^32^. To initiate this approach in *G. gynandra* we generated two F_2_ mapping populations in 2018 and 2019. Each was derived from founder lines of Malaysian and Malawian origin and assessed for vitamin content, and agriculturally important traits. Although photosynthesis has not been a major focus for crop improvement^33^, it has been noted that selecting solely for yield can overlook potential avenues for crop improvement such as photosynthetic enhancement^34^. Since the founders of Wag19 showed differences in traits underpinning the efficient C_4_ photosynthesis pathway^35^, we also assessed this population for variation in vein density, which is crucial for this pathway^36^. We anticipate that further similar analyses will enhance *G. gynandra* germplasm.

## Materials and Methods

### Production of plant material, growth and phenotyping

Two populations of *G. gynandra* were developed at Wageningen University & Research in 2018 and 2019, hereafter referred to as Wag18 and Wag19. For Wag18, the female and male parents of the F_1_ were derived from repeated self-fertilization of an early-flowering, short accession from Malaysia (TOT7200; Malaysia-03; MAY-03) and a late-flowering, tall accession from Malawi (TOT8917; Malawi-02; MAL-02; Sogbohossou et al., 2018). For Wag19, the female and male parents were derived from repeated self-fertilization of an early-flowering, short accession from Malaysia (TOT7199; Malaysia-01; MAY-01) and a late-flowering, short accession from Malawi (TOT8918; Malawi_01; MAL-01)^16^, these lines also have wide variation for C_4_ photosynthetic traits^35^. Wag18 consisted of 213 F_2_ lines and Wag19 187 F_2_ lines.

The populations were grown in the greenhouse of Wageningen University & Research, the Netherlands from March-June 2018 and April-July 2019. For Wag18 seven replicates of each parent were randomized, however, one replicate of Malaysia-03 was not healthy at the time of phenotyping and was therefore discarded. For Wag19 five replicates of each parent were randomized and two F_1_ lines were included. Growing conditions were tightly controlled with plants grown under irrigated conditions with temperatures maintained at 24^∘^C in the day and 20 ^∘^ C during night, and with artificial lights maintaining a minimum light intensity of 300 µmol m^−2^ s^−1^ and a photoperiod of 16hr days and 8hr nights. Sampling for DNA extraction and subsequent processing was carried out as described previously^29^.

For both Wag18 and Wag19, plant height was measured after 10 weeks and the leaf area of three fully mature leaves per plant was assessed using image analysis in ImageJ^37^. Flowering time was measured as the number of days from sowing to flowering. In Wag19, stem trichome density and stem colour were assessed as ordinal variables: 2 = dense trichomes or red colour; 1 = moderate trichome density and red/green colour; and 0 being no trichomes and green colour. Red colouration was indicative of anthocyanin content. Extraction and analysis of carotenoids and tocopherols as well as quantification was performed as described by^30^. In Wag18, seven plants died prior to vitamin-content assessment, along with an eighth plant that died before flowering. As a result, these plants were not assessed for plant height, leaf area and flowering time. Further, due to poor image quality or lack of replication, accurate leaf area could not be assessed for an additional nine plants. In Wag19, in total nineteen plants were not assessed for vitamin content due to sample contamination. Five plants died before being assessed for plant height, leaf area, stem colour, and stem trichome density. A further three died prior to flowering, and three plants could not be accurately assessed for leaf area. Lastly, six replicates of each parental line were assessed for vitamin content.

C_4_ leaf anatomical traits were assessed in Wag19 as follows. Plants were harvested over a three-day period 4 weeks after germination. The most recent fully expanded leaf was selected for sampling. Tissue was harvested from the central leaflet of each leaf and placed immediately in a plastic cuvette submerged in a 3:1 100% (v/v) ethanol:acetic acid fixative solution before treatment with 70% (v/v) ethanol solution overnight at 37^∘^C, being refreshed one hour into treatment. Samples were cleared using 5% (w/v) NaOH solution for two hours at 37^∘^C before being washed with, and replaced in, 70% (v/v) ethanol until preparation for imaging. Immediately prior to imaging, samples were immersed in Lugol’s solution (I_3_K), and washed with water to highlight vein tissue as described in Simpson et al. (2023). Samples were mounted with water and imaged on an Olympus BX41 light microscope with a mounted Micropuplisher 3.3 RTV camera (Q Imaging). Images were captured with Q-Capture Pro 7 software. Bundle sheath cells were imaged at x200 magnification and assessed using the line measurement tool in ImageJ^37^. Bundle sheath strand width was measured as the length of the bundle sheath strand perpendicular to the vein; six measurements were taken per field of view. Bundle sheath cell length was quantified by measuring the length parallel to the vein of 6 bundle sheath cells and dividing this by 6 to get a single value per field of view. For both parameters, six field of views were measured per plant. Four and five F_2_ lines were not assessed for bundle sheath cell width and length respectively, in addition to one replicate of Malaysia-01 due to poor clarity of microscopy images. The Wag19 lines were also assessed for vein density, using six fields of view per plant at a magnification of x100 using the Starch4Kranz pipeline^38^.

### SNP calling

To call and select Single Nucleotide Polymorphisms (SNPs) in the two sets of founder lines for genotyping-by-sequencing in each population, raw reads generated for each founder^29^ were checked for quality and trimmed using Trimmomatic^39^. The reads were aligned to the reference genome using the BWA MEM algorithm^40^ and Samtools^41^. Duplicate reads were filtered and read groups added using GATK (version 4.0) Variant calling and filtering was performed with the HaplotypeCaller and the VariantFiltration tool of GATK^42^. In order to select SNPs to be used for the characterization of the F_2_ population, heterozygous SNPs and SNPs with no call were discarded. Non-polymorphic SNPs between both parents were removed using vcffilterjdk and JEXL expressions^43^. SNP clusters in 150 bp windows were also removed. Furthermore, in the Wag18 founders, we subsequently used a set of re-sequenced genomes of 48 accessions: 24 from East and Southern Africa and 24 from Asia^29^ to select SNPs that were present in at least 12 accessions from the same region of origin as the parents and, we identified SNPs that were present in genomic regions of interest for our targeted traits irrespective of whether they were unique to the parental lines or shared by other accessions from the same region. These additional filtering steps were not applied to the Wag19 founders.

The final set consisted of 1309 SNPs and 9305 SNPs that were used for the genotyping-by-sequencing in the Wag18 and Wag19 population respectively.

### Heritably estimates, linkage maps, and QTL mapping

Broad-sensed Heritability (𝐻^2^) was measured according to Kearsey & Pooni^44^. The observed phenotypic variance 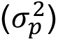 is the sum of the genetic variance 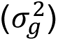 and environmental variance 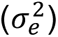. Since the homozygotic parental lines are genetically identical, the observed variance within each line is equivalent to 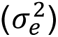 which can be extracted from the residual mean squares of the combined parental data. 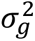 can therefore be measured with the following equation, where 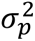 corresponds to the phenotypic variance excluding the parental lines:

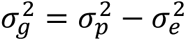

This can then be used to calculate broad sensed heritability:

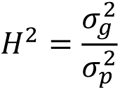

Heritability estimates were carried out by comparing parental phenotypes to that of the final mapping population.

In both Wag18 and Wag19 populations, reads were aligned to the reference genome^28^ using Bowtie2^45^ and Samtools^41^. Haplotype-based variant detection was carried out using Freebayes^46^ and filtering was undertaken with Bcftools^47^ before converting to R/qtl format using the R/utl package^48^. To produce the final SNP set, markers with a Minor Allele Frequency (MAF) less than 0.2 were removed along with any markers missing in more than 70% of individuals and ambiguous genotypes (i.e., being heterozygous in the parents or homozygous in the F_1_). This resulted in final SNP sets of 521 and 2035 for the Wag18 and Wag19 populations respectively.

Linkage Maps were generated using the following pipeline. First, duplicate individuals were identified. In Wag18, three pairs of duplicate individuals were identified that shared more than 95% of their genotypes; each pair was next to each other in the experimental design suggesting a sampling error. Due to not knowing which individual was correct all 6 individuals were removed. No duplicates were identified in Wag19. Secondly, we removed one individual in the Wag18 population that was completely homozygous. We then constructed genetic maps as detailed by Broman & Sen^49^ using the package R/qtl^50^. Marker positions were based on their physical location^28^, with errors being revealed through investigating marker recombination frequency, and successive use of the droponemarker function^49^. Local marker order was adjusted through rippling to minimize cross-over number by using a sliding window of 7 markers. Finally, individual genotyping errors were removed through study of double cross over rate and the calc.errorlod function. Our final map for Wag18 consisted of 206 individuals, 279 markers, with 99.9% genotype rate, and an error rate of less than 0.001 based on loglikelihood scores. For Wag19, we attained the same error rate, 187 individuals and 920 markers, with a 99.7% genotype rate.

QTL were detected in both populations using the Multiple QTL Mapping (MQM) method described by Broman & Sen^49^. First, single scans were performed using the scanone function. Second, two-dimensional scans were carried out using the scantwo function which considers QTL of large effect meaning modestly sized QTL can be identified in addition to interactions between QTL. Last, the putative QTL identified by scanone and scantwo scans were used as the starting model and the function stepwiseqtl used to identify the final QTL model for each trait. For all scans, Haley-Knott regression for interval mapping was implemented for QTL detection due to its rapidity and that our genotype datasets had very little missing data. Further, if single scans revealed significant QTL not identified through MQM, these were investigated for significance through linear regression. Significance thresholds were calculated based on 1000 permutations per trait per scan. QTL models were fitted from the output of stepwiseqtl and the applying the fitqtl function, so that Percentage of Variance Explained (PVE) for each QTL model could be calculated as:

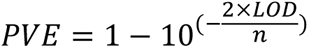

Where LOD is the logarithm of the odds score associated with the QTL or QTL model in question and n is the number of individuals phenotyped. Confidence intervals for each QTL were calculated as the 1.5 LOD drop intervals based on the final QTL model and scantwo permutations using the qtlStats function from R/qtlTools^51^. If the same QTL was identified for multiple traits, confidence limits were taken as being the overlapping region. Prior to mapping, phenotype data was transformed based on assessment of histograms in the mapping populations. Phenotype data were transformed when appropriate so that in the Wag18 population, leaf area and alpha-tocopherol-content underwent log-transformation, and in the Wag19 population, plant height was squared transformed, leaf area, and vein density were square-root transformed, flowering time was cube-root transformed, and alpha-tocopherol-content was log-transformed. To enable sufficient permutations to be generated for the non-parametric traits, stem trichome density and stem colour, normal models were selected while QTL mapping as it is accepted these still produce accurate results^49^, however single scans were compared to the non-parametric model to ensure QTL identified were correct. All QTL were identified under a significance threshold of 5% based on 1000 permutations derived from both the scanone and scantwo functions.

### Statistical analysis

All statistics was carried out in R (version 4.1.3). Variation between parent lines was analyzed using a two-sample Student t-test in R Studio (V: 4.0.0). Data were checked for normality and equal variance in each group using Shapiro-Wilks test and Bartlett’s test respectively. If data were not normal in one or both groups, the Mann-Whitney U (MWU) test was used in place of the Student’s t-test and variance was checked using Levene’s test. Results from Levene’s test determined comparisons made with the MWU test. If Levene’s test found equal variance, the MWU test for significance was based on median values. If Levene’s test found unequal variance between parental groups, the MWU test was based on variance. If data were normal in distribution but had unequal variance between groups, a Welch’s t-test was used. For mapping populations, normality tests were determined from data distribution, and pairwise Pearson correlation analysis carried out using R/GGally package^52^. To identify relationships between binary or discrete traits (stem trichome density, and stem colour) and continuous traits, ANOVA was used, while a Chi-squared test was used to test for a relationship between stem trichome density and stem colour. Linear regression was used to determine environmental effects and multiple QTL mapping performed as described. Plots were generated using R/Ggplot2^53^, R/corrplot^54^, R/qtlTools^51^, and R/qtl^50^. Logistic regression was carried out using the R/MASS package^55^, and p-values derived from resulting t-distributions by applying the cumulative distribution function.

## Results

### Phenotypic and heritability assessment for Wag18

Founders of the Wag18 population demonstrated statistically significant differences for all phenotypes assessed (𝑝 < 0.05, Table S1). With the exception of alpha-tocopherol, all other traits had higher values in Malawi-02 than Malaysia-03 (Figure 1). All traits displayed transgressive variation compared with the founders (Figure 1) and when broad-sense heritability (𝐻^2^) was calculated all were higher than 0.5 (Table S2), demonstrative of the fact they were grown under tightly controlled environmental conditions. Flowering time, plant height and leaf area correlated positively with each other, and the content of all three carotenoids were highly correlated (Figure 2).

**Figure 1:**
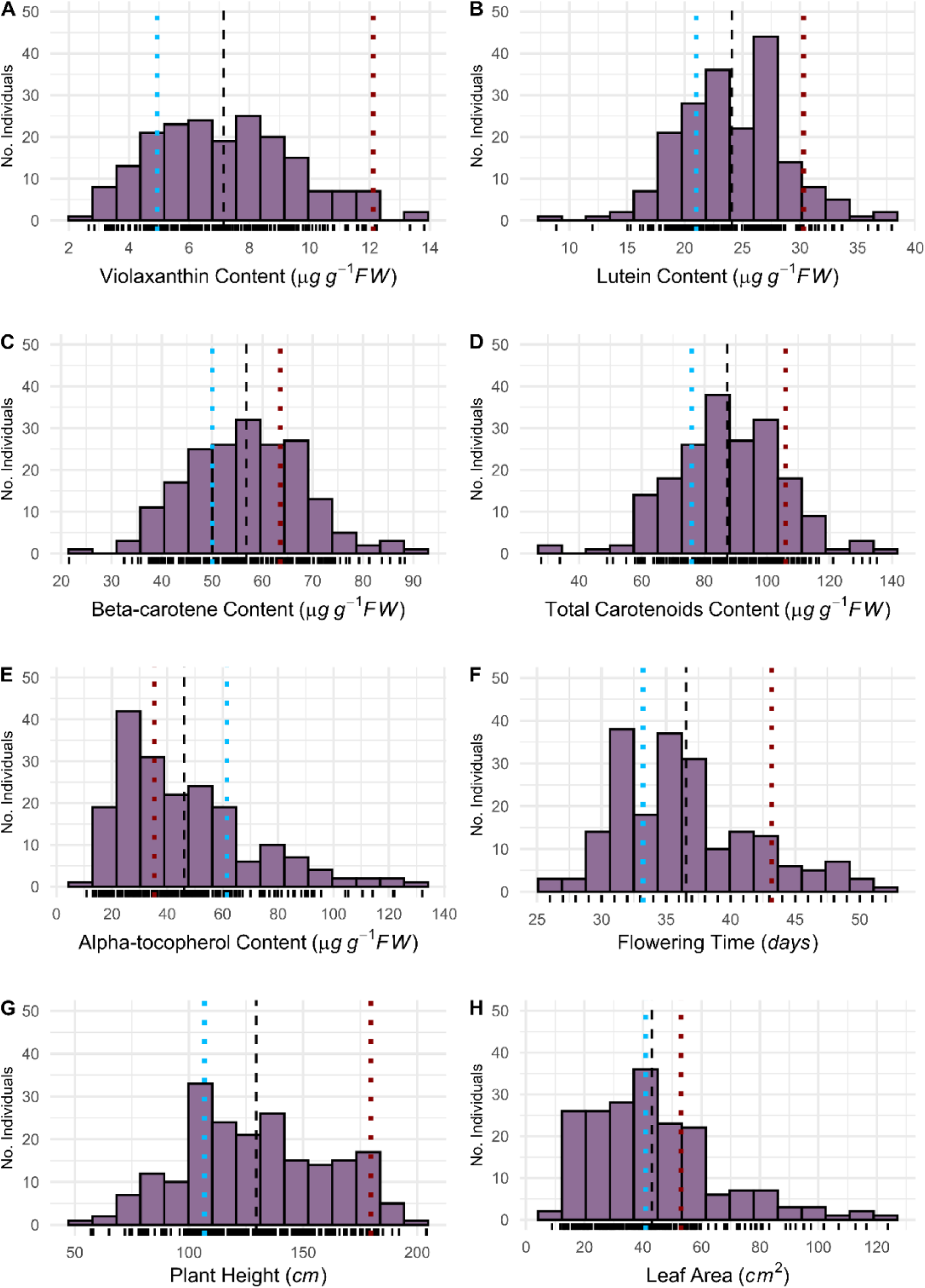
Transgressive variation for all traits in the Wag18 F_2_ mapping population. Blue and red dotted lines indicate means for the founders, Malaysia-03 and Malawi-02, respectively. The dashed black line represents is the mean of the F_2_ population. **(A)** Violaxanthin content; **(B)** Lutein content; **(C)** Beta-carotene content; **(D)** Total carotenoid content; **(E)** Alpha-tocopherol content; **(F)** Flowering time; **(G)** Plant height; **(H)** Leaf area.

**Figure 2:**
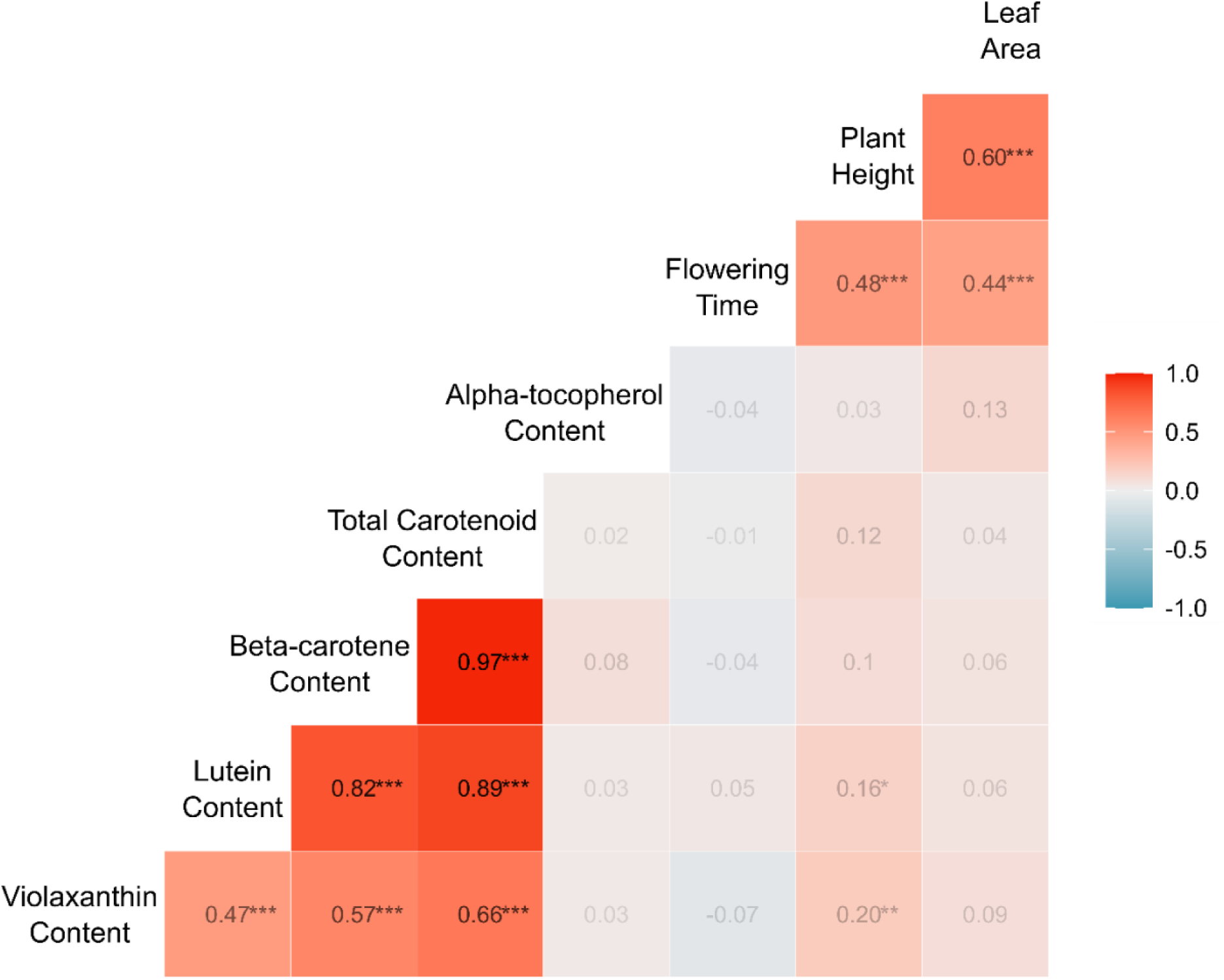
Correlation analysis for agronomic and nutrition traits in Wag18. Correlation coefficients were derived by Pearson correlation analysis and are presented in each box. Colour intensity indicates the strength of correlation, with deeper red representing a stronger positive relationship. The diagonal is the label for each phenotype. BSCW = Bundle Sheath Cell Width. Significance is shown as *** 𝒑 < 𝟎. 𝟎𝟎𝟏; ** 𝒑 < 𝟎. 𝟎𝟏; *𝒑 < 𝟎. 𝟎𝟓.

### Four unique QTL were identified in Wag18 across all traits

The Wag18 linkage map consisted of 279 markers covering 984.87cM with an average chromosome length of 82.07cM (Figure 3a & 3b). Across a population of 206 individuals, a total of 3293 cross over events were captured meaning an average recombination rate of 1.33 cross overs per individual per chromosome; we attained a genotyping error rate of approximately 0.

**Figure 3:**
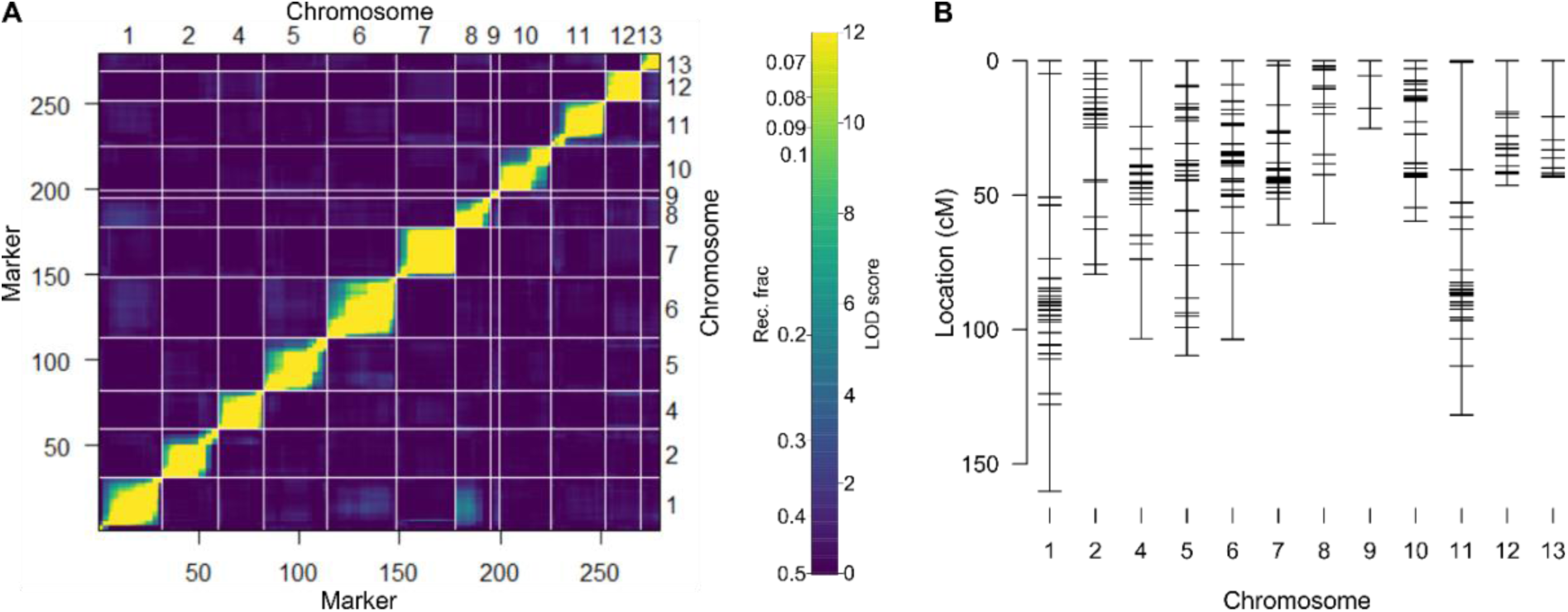
Wag18 genetic map. **(A)** Heatmap displaying recombination frequencies and LOD scores between each marker in the upper and lower triangles respectively; with yellow indicating a lower recombination frequency / higher LOD score, hence suggesting marker-pairs are linked. **(B)** Final genetic map.

In Wag18, a total of seven putative QTL (pQTL) were identified across all traits (significance less than 0.05 based on 1000 permutations, Table S3). The two pQTL identified for violaxanthin content, 1:84.7 and 1:89.8 were very close together yet had opposite additive effects (Table S3). The nearest markers for these two QTL were 1.05951559 (located on chromosome 1 at position 5951559bp) and 1.13366757 respectively, for both markers, the AB genotype had significantly higher violaxanthin content than the AA (Malaysia-02) genotype (𝑝 = 0.033; 𝑡 = −2.15; 𝑑𝑓 = 132.55 and 𝑝 < 0.001; 𝑡 = −3.43; 𝑑𝑓 = 121.43 respectively). However, it appears that the overall negative additive effect of 1:84.7 was an artifact of a small number of individuals with the Malawi-02 genotype (𝑛 = 20) and so this pQTL was not pursued. The remaining violaxanthin pQTL (1:89.8) was not deemed significant through multiple QTL mapping, nor was it identified in single QTL scans so was also not pursued. A pQTL identified for plant height (1:90.8) and leaf area (1:88.9) had similar positive additive effects resulting from the Malawi-03 (B) allele (Table 1) while also being situated in the same region, as these were strongly correlated traits (Figure 3), these pQTL were considered as the same unique QTL controlling size, hereafter w18Size1.

**Table 1:** Overall summary of QTL identified in the Wag18 population.

| Phenotype | QTL number | QTL name | Genetic map position (chr:cM) | Nearest marker (chr.bp) | Physical position (Chr:Mb) | Physical range (Mb) | Additive effect | Dominance effect | LOD | Model formula | Model LOD | PVE by model (%) |
| --- | --- | --- | --- | --- | --- | --- | --- | --- | --- | --- | --- | --- |
| Lutein content | Q1 | w18L1 | 2:18.0 | 2.2745371 | 2:9.34-27.87 | 18.53 | -2.08 | 0.01 | 4.64 | y ~ Q1 | 4.64 | 10.23 |
| Alpha-tocopherol content* | Q1 | w18aT1 | 9:5.7 | 9.00780456 | 9:0.00-1.32 | 1.32 | -0.42 | 0.10 | 17.41 | y ~ Q1 | 17.41 | 33.16 |
| Flowering time | Q1 | w18FT1 | 6:45.0 | 6.36025417 | 6:27.36-44.29 | 16.93 | 3.00 | -2.02 | 9.19 | y ~ Q1 | 9.19 | 19.24 |
| Plant height | Q1 | w18Size1 | 1:90.8 | 1.13687113 | 1:11.74-20.24 | 8.50 | 25.12 | -1.30 | 12.38 | y ~ Q1 | 12.38 | 25.02 |
| Leaf area* | Q1 | w18Size1 | 1:88.9 | 1.11928029 | 1:11.74-20.24 | 8.50 | 0.38 | -0.11 | 8.33 | y ~ Q1 | 8.33 | 18.38 |

*PVE = Percentage of Variance Explained, QTL name is based on population and phenotypic information. Genetic map position is shown as "Chromosome:position (cM)", nearest marker name is based on physical position: "Chromosome.position (bp)". Negative and positive additive effects mean the allele from Malaysia-03 and Malawi-02 respectively is responsible for an increase in the trait. \*Analysis carried out on log transformed data. Numbers in italics were derived non-parametrically.*

To summarise, extensive variation and high heritabilites were recorded for Wag18, and this enabled the mapping of four unique QTL associated with six agriculturally important traits (Table 1). w18Size1 explained 25% and 18% of the phenotypic variance observed for plant height and leaf area respectively; w18L1 explained 10% of the lutein content variance; w18aT1 contributed to 33% of the variance seen for alpha-tocopherol content; w18FT1 explained 19% of the observed flowering time.

### Phenotypic relationships in the Wag19 population

When the Wag19 founders were assessed Malawi-01 had higher values than Malaysia-01 for carotenoid content, flowering time, plant height and leaf area, and a lower alpha-tocopherol content (Figure 4), but the difference observed for all carotenoids and plant height was not statistically significant (Table S3). For traits measured in Wag19 that were not assessed in Wag18, compared with Malaysia-01, vein density, trichome density, and stem colour was significantly higher in the Malawi-01 parent, while bundle sheath strand width and cell length were lower (Table S3). Traits in Wag19 displayed transgressive segregation, except bundle sheath strand width and vein density, for which Malawi-02 demonstrated extremely high and low values respectively (Figure 4).

**Figure 4:**
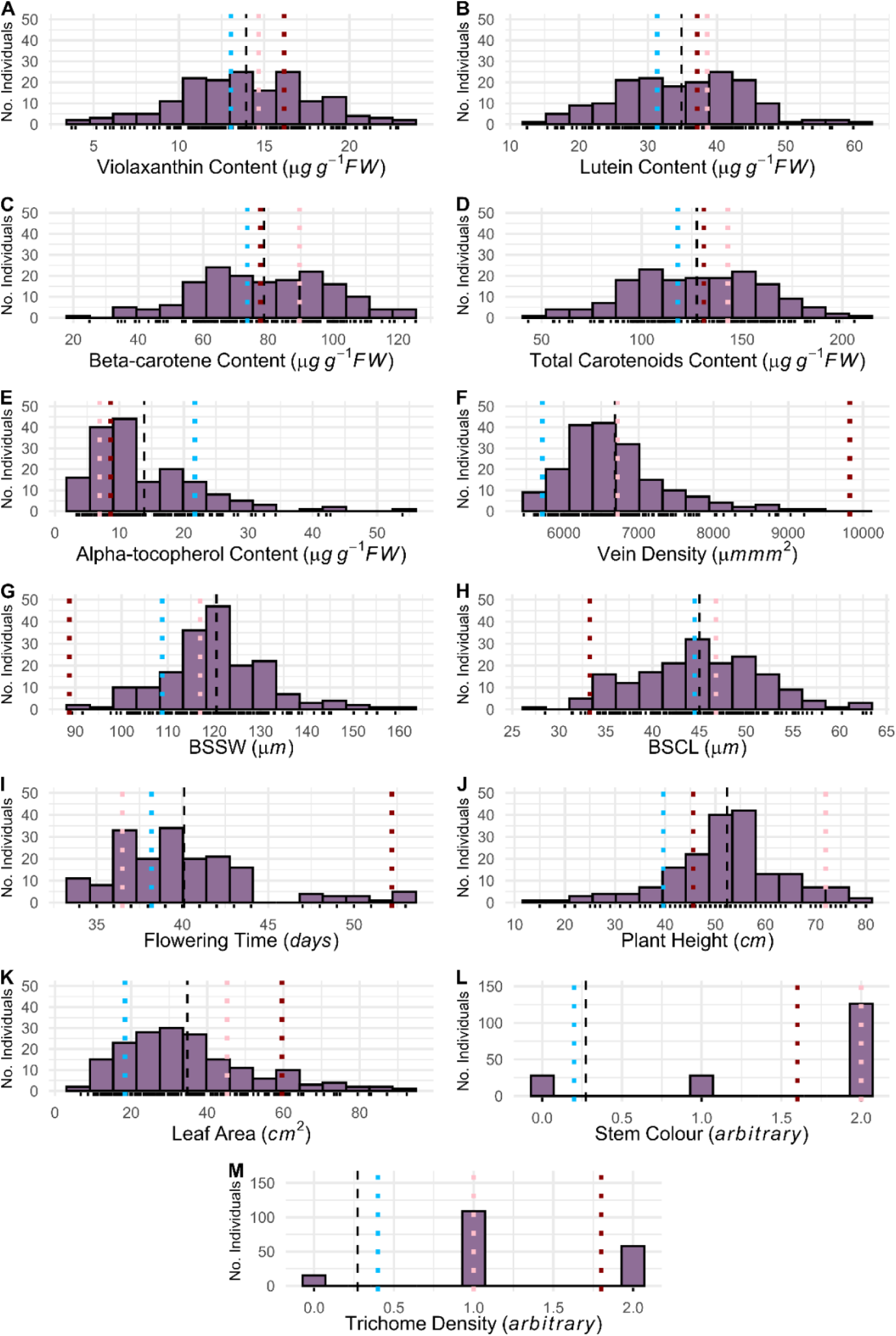
Phenotypic distributions in the Wag19 F_2_ mapping population. The blue, red and purple dotted lines indicate the means for Malaysia-01, Malawi-02, and the F_1_s respectively. The dashed black line represents the mean of the F_2_ mapping population. **(A)** Violaxanthin content; **(B)** Lutein content; **(C)** Beta-carotene content; **(D)** Total carotenoids content; **(E)** Alpha-tocopherol content; **(F)** Vein density; **(G)** Bundle Sheath Strand Width (BSSW); **(H)** Bundle Sheath Cell Length (BSCL); **(I)** Flowering time; **(J)** Plant height; **(K)** Leaf area; **(L)** Stem colour; **(M)** Trichome density.

Heritabilities were generally lower in the Wag19 population compared to Wag18, however only plant height (0.49) and bundle sheath cell length (0.19) had heritability of less than 0.5 (Table S2). Flowering time correlated with leaf area in Wag19, but this was not the case for plant height (Figure 5). Rather, in Wag19, plant height was also positively correlated with leaf area. Analysis of variance found that flowering time differed between stem colour categories (𝑝 < 0.001, 𝐹 = 10.07, 𝑑𝑓 = 178), with stems that lacked anthocyanin flowering earlier. As in Wag18, carotenoid content of the three carotenoids tested were highly correlated, while alpha-tocopherol content was significantly correlated with total carotenoid content, beta-carotene, and violaxanthin content (Figure 4). Traits related to C_4_ photosynthesis such as vein density, bundle sheath strand width, and bundle sheath cell length were strongly correlated, with the former having a negative relationship with the latter two traits. Interestingly, bundle sheath strand width was slightly, albeit significantly negatively correlated with carotenoid content. Plant height was positively correlated with bundle sheath size but negatively with vein density. A Chi-squared test revealed no significant relationship between stem colour and stem trichome density.

**Figure 5:**
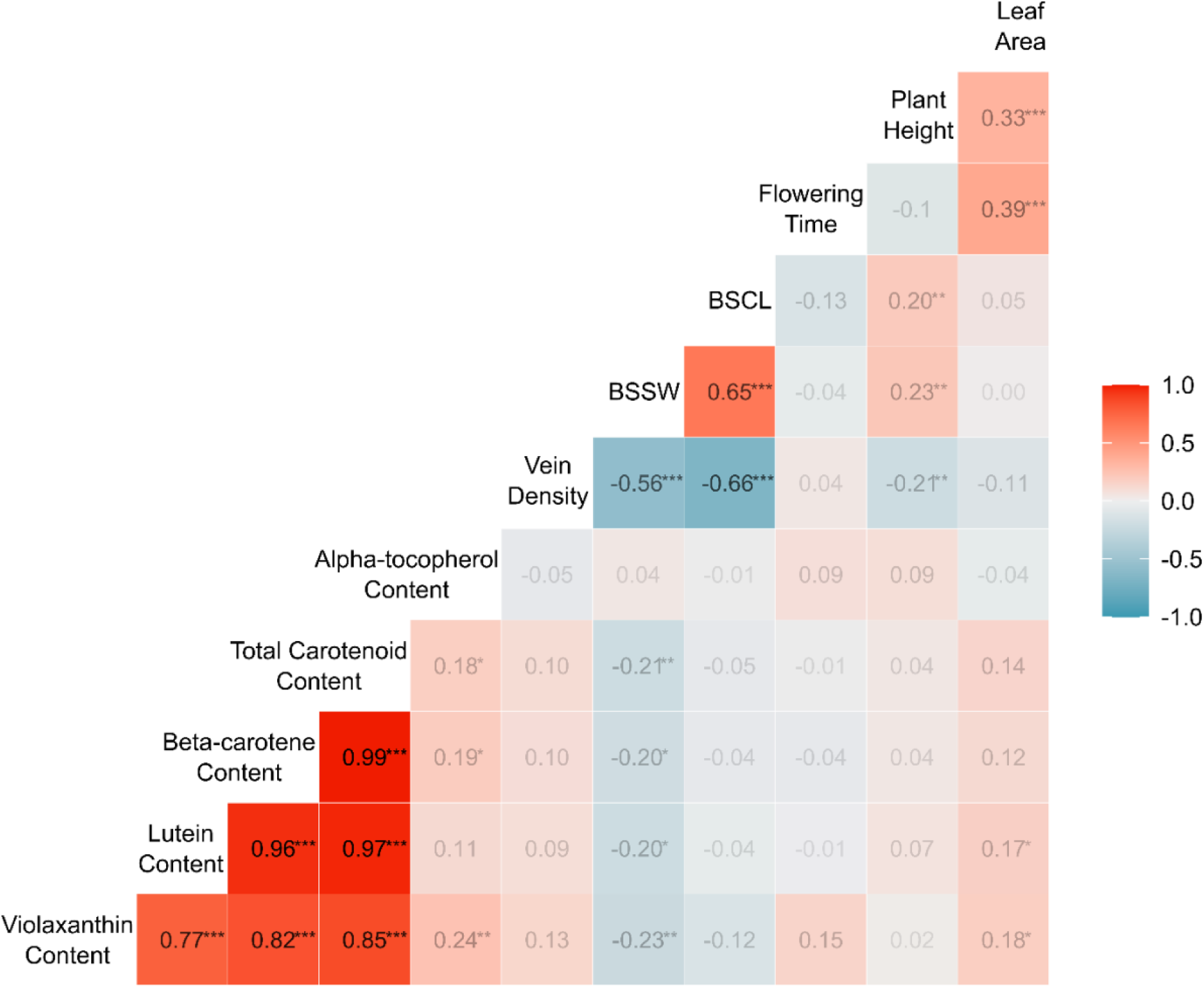
Correlation analysis for agronomic, nutrition and C_4_ traits in the Wag19 population. Numbers indicate correlation coefficients derived by Pearson correlation analysis. Colour intensity indicates the strength of correlation, with deeper red representing a stronger positive relationship. The diagonal is the label for each phenotype. BSSW = Bundle Sheath Strand Width, BSCL = Bundle Sheath Cell Length (measured as length of 6 cells). Significance is shown as *** 𝒑 < 𝟎. 𝟎𝟎𝟏; ** 𝒑 < 𝟎. 𝟎𝟏; *𝒑 < 𝟎. 𝟎𝟓.

### Increased marker density in Wag19 captured more cross-over events compared with Wag18

The Wag19 linkage map consisted of 920 markers covering 1645.02cM with an average chromosome length of 126.54cM (Figure 6a & 6b). In a population of 187 individuals, a total of 5465 cross over events were captured meaning an average recombination rate of 2.35cross overs per individual per chromosome. Both linkage maps had a genotyping error rate of approximately 0. While the smallest four chromosomes, 3, 15, 16, and 17 (Hoang *et al.,* 2023) remained unmapped, the improved marker availability in Wag19, due to a less stringent selection process compared to Wag18 meant that chromosome 14 was captured in Wag19.

**Figure 6:**
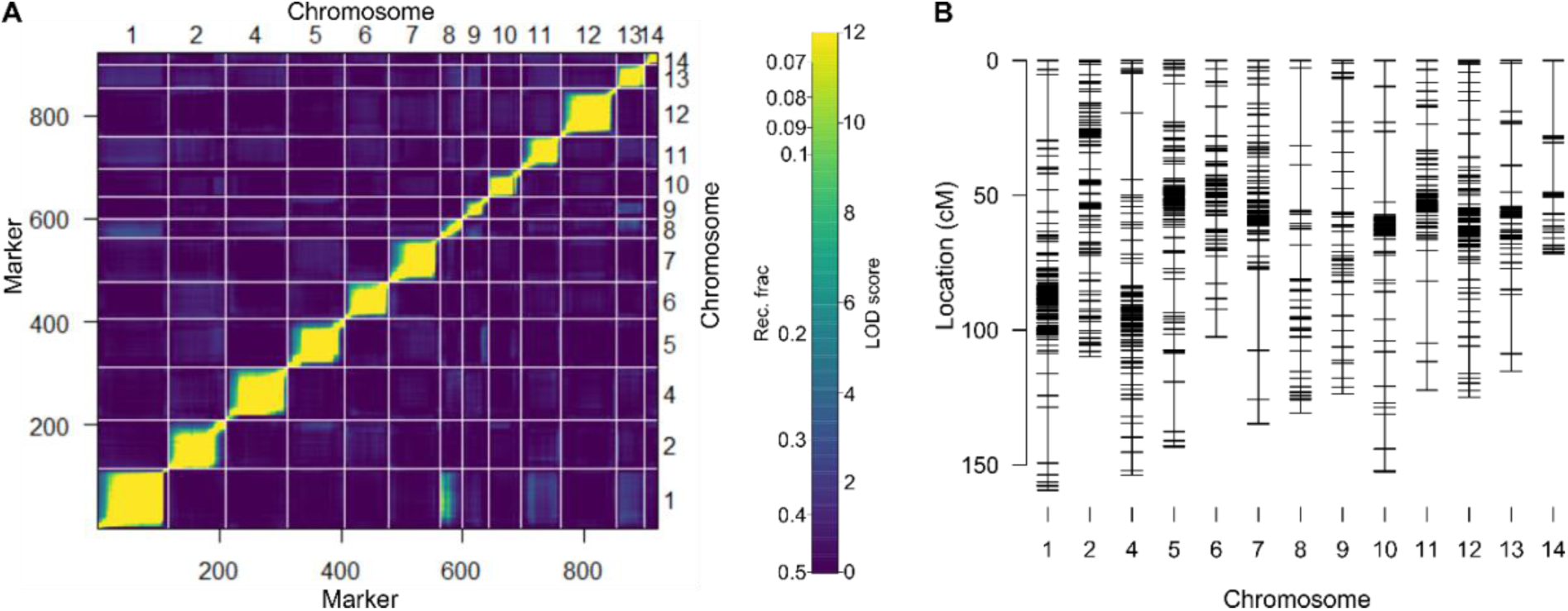
Wag19 genetic map. **(A)** Heatmap displaying recombination frequencies and LOD scores between each marker in the upper and lower triangles respectively; with yellow indicating a lower recombination frequency / higher LOD score, hence suggesting marker-pairs are linked. **(B)** Final genetic map.

### Eleven unique QTL were initially identified in the Wag19 population

In Wag19, a total of fourteen pQTL were identified (significance less than 0.05 based on 1000 permutations, Table S5). Two pQTL were found for vein density, one on chromosome 4 at position 122.00cM (4:122.0; 𝐿𝑂𝐷 = 3.88) and one on chromosome 14 at position 6.00cM (14:6.0; 𝐿𝑂𝐷 = 4.30; Figure S1a). Looking at the nearest markers for these two traits (4.59646348 and 14.00360911) it was clear the increase in vein density was, as expected, driven by the Malawi-01 (B) allele (Figure S1b & S1c), however 14.00360911 had just a single individual that was homozygous for the B allele, when this line was ignored, linear regression of the genotypes at these two markers on vein density found only 4:122.0 to be significant (𝑝 < 0.001), hence we considered this to be the only true QTL associated with vein density and hereafter refer to it as w19VD1.

pQTL for plant height (1:83.5), and leaf area (1:84.3) were found at approximately the same location and were positively influenced by the Malawi-01 (B) allele. Since these traits are also significantly correlated (𝑟 = 0.33; 𝑝 < 0.001, Figure 5), we consider these pQTL to represent the same multi-trait QTL, which appears to affect overall plant size; labelled w19Size1. Further, we identified two pQTL each for both discrete traits, stem trichomes (7:17.9 and 8:129.0) and stem colour (6:51.0 and 7:3.9), including a significant interacting term between the latter two. Since these were discovered under the assumption of normality, derived models (Table S5) were checked using ordinal regression (Table S6). This found that for stem trichomes the optimal model (𝐴𝐼𝐶 = 274.65) included both 7:17.9 and 8:129.0 having significant effects. While for stem colour, the optimal model (𝐴𝐼𝐶 = 160.91) included both 6:51.0 and 7:3.9, the pQTL 7:3.9 was not significant. Considering the minimal model including just 6:51.0 had only a slightly higher AIC (𝐴𝐼𝐶 = 160.91), we do not currently consider 7:3.9 a high priority QTL to pursue; including the interacting term had no effect on this analysis. Additionally, the single scans derived parametrically and non-parametrically were very similar for stem trichome density (Figure S2a) however for stem colour, the LOD peak was smaller using the non-parametric method (Figure S2b). To maintain accuracy, final LOD scores for QTL associated with these traits were based on non-parametric single scans.

In summary, eleven unique QTL affecting eight traits were identified in Wag19 (Table 2). One QTL was found to be significant for violaxanthin content (w19V1); one QTL for alpha-tocopherol content (w19aT); one for vein density (w19VD1); one QTL for flowering time (w19Ft1), one unique QTL for plant height (w19H1); two QTLs for stem trichomes (w19ST1 and w19ST2); one QTL for stem colour (w19SC1); and one QTL was found to effect overall plant size, i.e., having the same effect on plant height and leaf area (w19Size1). Overall, 12% of the variance observed for violaxanthin content, 14% for alpha-tocopherol content, 8% for vein density, 24% for flowering time, 25% for plant height, 29% for leaf area, 21% for stem trichomes and 52% for stem colour was explained by these QTL (Table 2). For traits assessed in both populations, four QTL were mapped in Wag18 (Figure 7a) and seven were mapped in Wag19 (Figure 7b).

**Figure 7:**
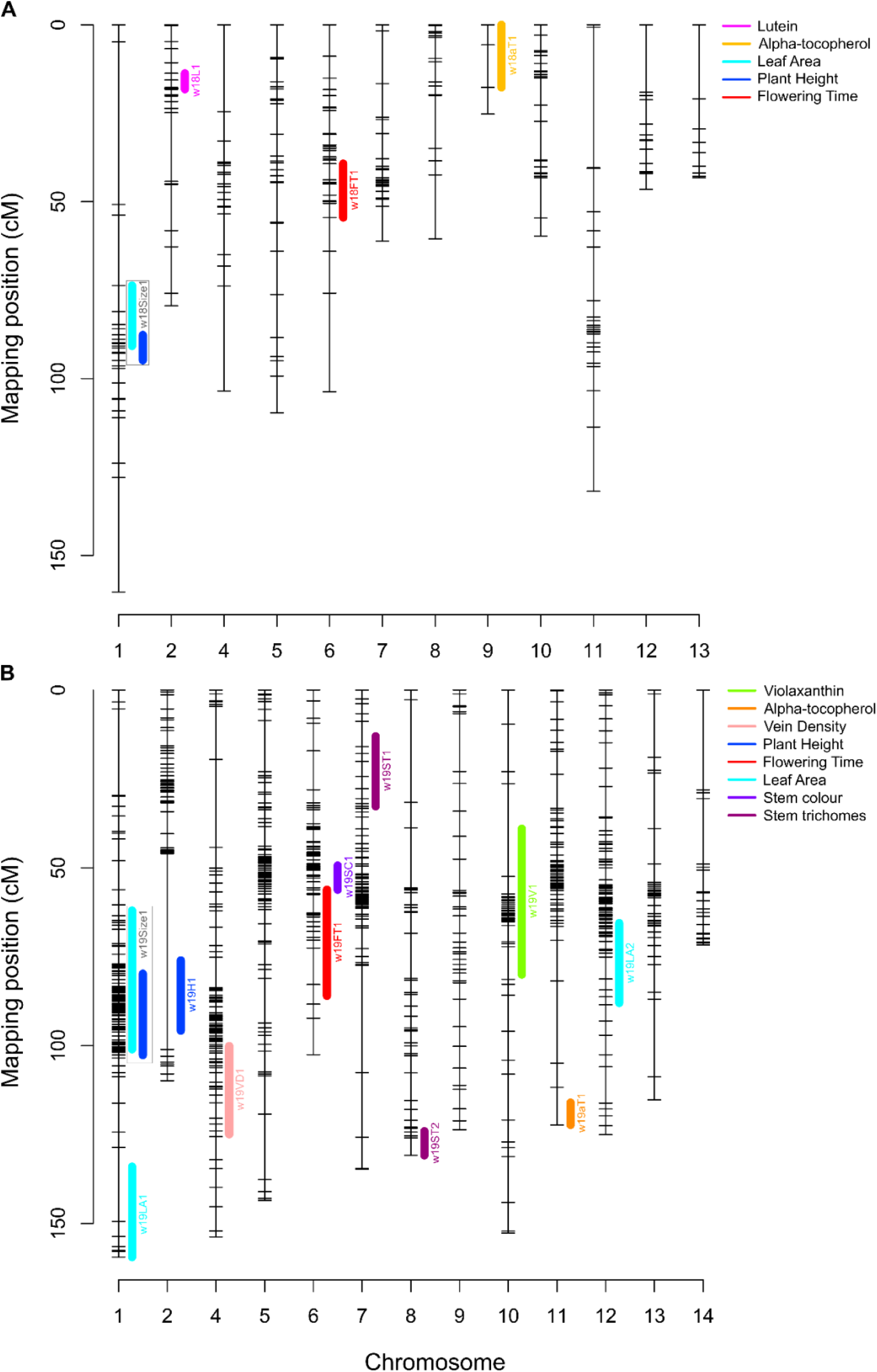
Location of QTL in the (A) Wag18 and (B) Wag19 populations. QTL associated with each trait are colour-coded. Multi-trait QTL are represented in grey boxes. Confidence intervals were identified using the R/qtlTools package (Lovell, 2021).

**Table 2:** Overall summary of QTL identified in the Wag19 population.

| Phenotype | QTL number | QTL name | Genetic map position (chr:cM) | Nearest marker (chr.bp) | Physical location (Chr:Mb) | Physical range (Mb) | Additive effect | Dominance effect | LOD | Model formula | Model LOD | PVE by model (%) |
| --- | --- | --- | --- | --- | --- | --- | --- | --- | --- | --- | --- | --- |
| Violaxanthin content | Q1 | w19V1 | 10:58.4 | 10.03027582 | 10:0.70-39.26 | 38.56 | 1.07 | -2.20 | 4.77 | $y \sim Q1$ | 4.77 | 12.26 |
| Alpha-tocopherol content* | Q1 | w19aT1 | 11:122.3 | 11.41198362 | 11:4.09-41.20 | 37.11 | -0.23 | -0.20 | 5.56 | $y \sim Q1$ | 5.56 | 14.14 |
| Vein Density**** | Q1 | w19VD1 | 4:122.0 | 4.59646348 | 4:53.59-60.56 | 6.97 | 1.40 | -0.36 | 3.88 | $y \sim Q1$ | 3.88 | 8.75 |
| Flowering time** | Q1 | w19FT1 | 6:77.0 | 6.46920704 | 6:36.68-49.41 | 12.73 | 0.083 | -0.033 | 10.85 | $y \sim Q1$ | 10.85 | 24.35 |
| Plant height*** | Q1 | w19Size1 | 1:83.5 | 1.17961377 | 1:11.15-61.79 | 50.64 | 514.58 | 419.87 | 7.46 | $y \sim Q1 + Q2$ | 11.62 | 25.47 |
| Plant height*** | Q2 | w19H1 | 2:84.0 | 2.38898522 | 2:38.62-39.24 | 0.63 | -528.14 | 12.25 | 4.48 | $y \sim Q1 + Q2$ | 11.62 | 25.47 |
| Leaf area**** | Q1 | w19LA1 | 1:145.0 | 1.70602349 | 1:70.06-70.92 | 0.85 | -0.27 | 0.99 | 4.98 | $y \sim Q1 + Q2 + Q3$ | 14.51 | 31.15 |
| Leaf area**** | Q2 | w19Size1 | 1@84.3 | 1.18700207 | 1:11.15-61.79 | 50.64 | 0.53 | 0.28 | 4.59 | $y \sim Q1 + Q2 + Q3$ | 14.51 | 31.15 |
| Leaf area**** | Q3 | w19LA2 | 12@73.0 | 12.38068542 | 12:2.70-39.64 | 36.94 | 0.73 | -0.19 | 5.75 | $y \sim Q1 + Q2 + Q3$ | 14.51 | 31.15 |
| Stem trichomes | Q1 | w19ST1 | 7:17.9 | 7.00721601 | 7:0.65-1.29 | 0.64 | -0.26 | -0.08 | 4.63 | $y \sim Q1 + Q2$ | 8.87 | 20.10 |
| Stem trichomes | Q2 | w19ST2 | 8:129.0 | 8.44525256 | 8:44.28-44.53 | 0.25 | 0.25 | 0.15 | 4.24 | y ~ Q1 + Q2 | 8.87 | 20.10 |
| Stem colour | Q1 | w19SC1 | 6:51.0 | 6.30828142 | 6:23.19-36.67 | 13.48 | 0.73 | 0.76 | 29.37 | y ~ Q1 | 29.37 | 52.44 |

*PVE = Percentage of Variance Explained; nearest marker name is based on physical position: "Chromosome.position(bp)". Negative and positive additive effects mean the allele from Malaysia-01 and Malawi-01 respectively is responsible for an increase in the trait. Analysis carried out on \*log, \*\*cube-root, \*\*\*squared, \*\*\*\*square-root transformed data. Numbers in italics were derived non-parametrically.*

### Agriculturally important QTL found in both populations

In Wag18, the QTL w18Size1 was located on chromosome 1 with a peak around 89.9cM and in Wag19 the QTL w19Size1 was located on chromosome 1 with a peak around 85.9cM (Figure 7). The nearest marker to both these QTLs were 1.13366757 and 1.17199903 in Wag18 and Wag19 respectively and in both cases, the “African” allele (B) coming from Malawi-02 in Wag18 and Malawi-01 in Wag19, was responsible for the increase in plant size (Figure 8a & 8b). The confidence limits based on the overlapping regions of the traits they were associated with (Figure 7) for these QTL was between 87.7cM and 90.7cM for w18Size1 and between 79.7cM and 101.0cM for w19Size1. Based on the nearest markers, the corresponding physical range for these QTL was 11.74-20.24Mb and 11.15-61.79Mb. While this region contains many genes, the overlapping QTL region means these effects may be under the control of the same gene or group of genes^56^.

**Figure 8:**
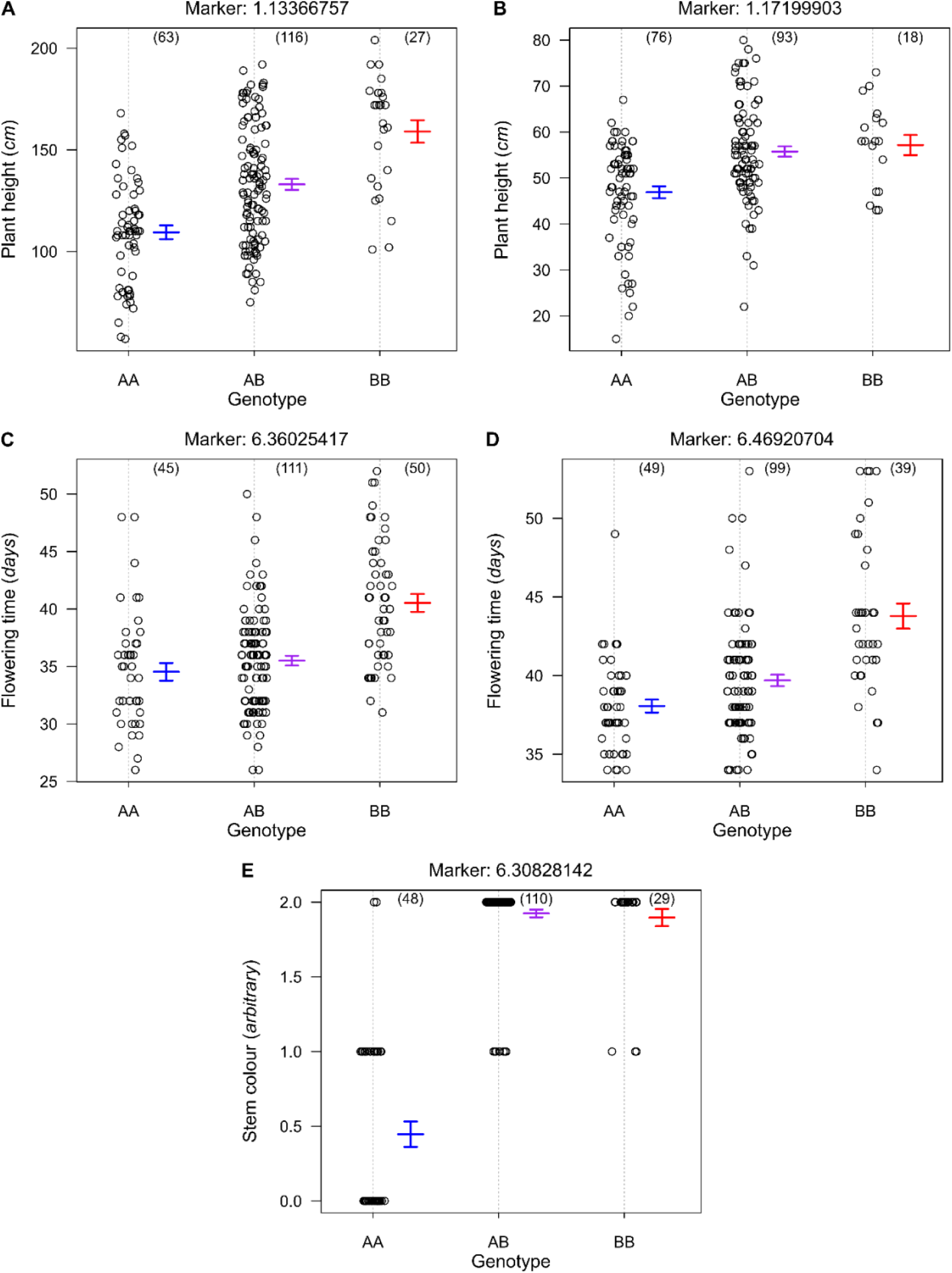
Phenotypes for agronomically important traits plotted against genotypes at markers nearest to associated QTL. **(A)** Plant height (indicative of plant size) increases at the marker nearest to w18Size1 when plants have the “African” allele; **(B)** the same trend is observed for w19Size1. **(C)** Flowering time increases when plants have the B allele in the marker closest to w18Ft1 and **(D**) w19Ft1. **(E)** All individuals homozygous for the Malaysia-01 (A) allele had no visible anthocyanin. Numbers in parentheses are the number of individuals with each genotype. Confidence intervals for average phenotypes within each genotype group are shown as blue for AA, purple for AB, and red for BB.

Further, the QTL w18Ft1 was located on chromosome 6 with a peak around 45.0cM and w19Ft1 was on chromosome 6 with a peak around 77.0cM (Figure 7). Once again, the African allele was responsible for the later flowering time (Figure 8c & 8d). The confidence limits for these QTL were 39.2 - 54.5cM and 54.1 - 86.0cM corresponding to a physical range of 27.36 - 44.29Mb and 36.68 - 49.41Mb for w18Ft1 and w19Ft1 respectively. This overlap suggests the same underlying gene or group of genes may be responsible for this effect.

## Discussion

### Stable QTL provide clear avenues for prebreeding in *G. gynandra*

We identified significant QTL in *G. gynandra*, which can be integrated into targeted breeding programs. This advancement paves the way for the widespread adoption of *G. gynandra* as a food crop. Given that *G. gynandra* is currently harvested as a wild plant with minimal domestication^57^, we anticipate these QTL will contribute to the development of improved germplasm, enhancing its potential for breeding efforts.

Two pairs of QTL were identified, w18Size1 and w19Size1 and w18Ft1 and w19Ft1, that mapped to the same regions in both populations and could be the result of the same underlying gene(s; Lynch and Walsh, 1998). These controlled plant size and flowering time (Figure 7). Interestingly, their effects were relatively stable in both populations, in the Wag18 population, w18Size1 explained 25% and 18% of the variance associated with plant height and leaf area respectively, which was similar to the variance explained by w19Size1 for these two traits, 17% and 10%, especially when considering the lower heritabilities associated with the Wag19 population (Table S2). Further, the w18Ft1 and w19Ft1 QTL explained 19% and 24% of the variance associated with flowering time in each population. Considering that *G. gynandra* is a commercially viable crop with nutritional and medical applications^16,18–21^, identifying QTL associated with flowering time and plant size, two traits of value to plant breeders, in multiple environments and populations, and given their physical locations are known (Table 1 & 2), means markers could be generated so desirable alleles could be incorporated into breeding germplasm via marker assisted selection^58^. To our knowledge, these are the first example of stable targets for marker assisted selection identified in *G. gynandra* and could be of great benefit in the next stages of this crop’s transcendence from orphan to viable.

Further, across both populations, we identified four QTL associated with the content of the carotenoids, lutein and violaxanthin, and the vitamin E, alpha-tocopherol. Anthocyanin content was assessed ordinally and only in the Wag19 population, nonetheless it’s associated QTL, w19SC1, explained 52.44% of total variance associated with stem colour (Table 2). Close inspection of the nearest marker to this QTL located on chromosome 6, 6.30828142, found just 2 out of 48 AA (Malaysia-01 genotype) individuals had a score of 2 (indicative of high anthocyanin) and no individuals with even one copy of the B (Malawi-01) allele were absent anthocyanin (a score of 0; Figure 7e). Further, w19SC1 was located in close proximity to w19Ft1, controlling flowering time, on chromosome 6 (Figure 6b), for which also the Malaysia-03 genotype had the earlier flowering time. Since ANOVA found stems lacking anthocyanin flowering significantly earlier (𝑝 < 0.001, 𝐹 = 10.07, 𝑑𝑓 = 178), it is possible these two QTL are linked or controlled by the same underlying gene(s). Further, given this relationship, and that the occurrence of anthocyanin is present from an early stage of plant development, it could serve as a useful phenotypic marker for agricultural planning.

The vitamins we assessed are involved in protection of eyes from diseases, age-related degeneration^59,60^, mitigating UV-induced skin damage^61^ and are important for the immune system^62^. There is a need for a targeted crop improvement programme for *G. gynandra*^16,57^ and these identified QTL could be useful in marker assisted selection for vitamin content. Furthermore, a previously reported negative correlation between carotenoids and tocopherols^30^ was not identified in the Wag18 population (Figure 2), and was observed only weakly in the Wag19 population (Figure 5) suggesting these two traits, can in fact be bred for simultaneously. A brief assessment of the F_1_ lines in the Wag19 population also found strong heterosis for plant height (Figure 4j). This has not been reported previously for *G. gynandra* and indicates its suitability for hybrid breeding^63^, which is of huge benefit for generating advanced breeding programmes^15^.

### Linkage mapping can identify QTL associated with the C_4_ syndrome

Cleomaceae is the closest related family to the Brassicaceae and therefore contains the most closely related C_4_ species to the model C_3_ plant *Arabidopsis thaliana* with which *G. gynandra* shares high similarity in gene sequence^64^. This makes it an excellent model organism for the study of C_4_ photosynthesis and it has been used as a system through which genetic determinants of C_4_ enzymes have been investigated^23–27^. C_4_ photosynthesis has evolved independently over 65 times^65^ and is more energy efficient under warm, dry conditions compared to the C_3_ state. Mathematical modelling suggests that morphological changes such as increased vein density and larger bundle sheath cell size were crucial in the early evolution of the C_4_ syndrome^36^.

Kranz traits, vein density and bundle sheath size showed extensive variation (Figure 4f, 4g & 4h) with one QTL, w19VD1 being mapped to chromosome 4 (Figure 7b). Further, initial QTL scans found an additional pQTL for vein density (Table S4). Although we deprioritized 14:6.0 as an outlier appeared to skew the results, rendering this pQTL association statistically insignificant after its removal, it is possible that this represents a genuine association. This is especially considering the additive effect observed in the heterozygote, and the large vein densities found in both ‘BB’ lines assessed (the Malawi-01 founder line and the F_2_ line in question; Figure 1f & S1c). A population with a greater prevalence of these alleles would be needed to demonstrate this statistically.

Furthermore, it should be noted that in *G. gynandra* and other C_4_ plants, carotenoids and tocopherols are crucial for protecting the photosynthetic apparatus from oxidative stress, ensuring efficient photosynthesis under intense light conditions and other abiotic stressors such as drought and high temperature^66,67^, therefore the aforementioned QTL linked to these traits could also be useful for identifying lines suitable for growing under more extreme conditions.

To our knowledge, this is only the first example of linkage mapping being employed to identify QTL associated with C_4_ characteristics in any dicotyledon. In maize (*Zea mays*) a bi-parental population identified QTL for stomatal size and density^68^, and so together these data provide strong support for the notion that quantitative genetics could be used to study C_4_ traits^69^. Given the region containing w19VD1 contains 318 genes^28^, we have refrained from reporting candidate genes due to the ambiguity in searching for GO terms amongst a relatively large number of genes. Instead, we aim here to prove that reverse genetics approaches such as QTL mapping is a viable option for the study of C_4_ photosynthesis. Furthermore, alongside a wild diversity panel, there is currently a Multi-Parental Advanced Generation Inter-Cross (MAGIC) population of *G. gynandra* being developed with the aim of mapping traits related to C_4_ photosynthesis. Such association mapping methods are prone to false positives^70^ so populations derived from simple bi-parental crosses, such as that assessed here, that are equally related and therefore absent population structure, can add power to and complement such studies that enable higher resolution and the fine mapping of causal genes^58,69^. We demonstrate here the effectiveness of mapping QTL in *G. gynandra* and, given its rapid generation time, phenotypic and genotypic diversity^30,35^, combined with the power of bi-parental populations^58^, it is an excellent resource for further mapping of additional traits. Furthermore, many F_1_ hybrids already exist that can be used as a resource for future study of developmental, physiological or agricultural traits in *G. gynandra*.

## Author contributions

CJCS, PS, DS, ES, and JMH conceptualized and designed the study. CJCS, DS, and PS collected data. DS performed extracts and made the crosses from which Wag18 was derived. GR made the crosses from which Wag19 was derived. DS called SNPs for genotyping-by-sequencing. CJCS mapped SNPs to the reference genome. CJCS carried out microscopy, heritability and QTL analysis. CJCS generated the figures. CJCS, PS, JMH, and ES wrote the article. JMH, PS and ES oversaw and supervised this work.

## Data availability

Genotype and phenotype data can be found at https://github.com/plycs5/GgQTL. Sequencing datasets are available on request.^1^

---

**Figure S1:**
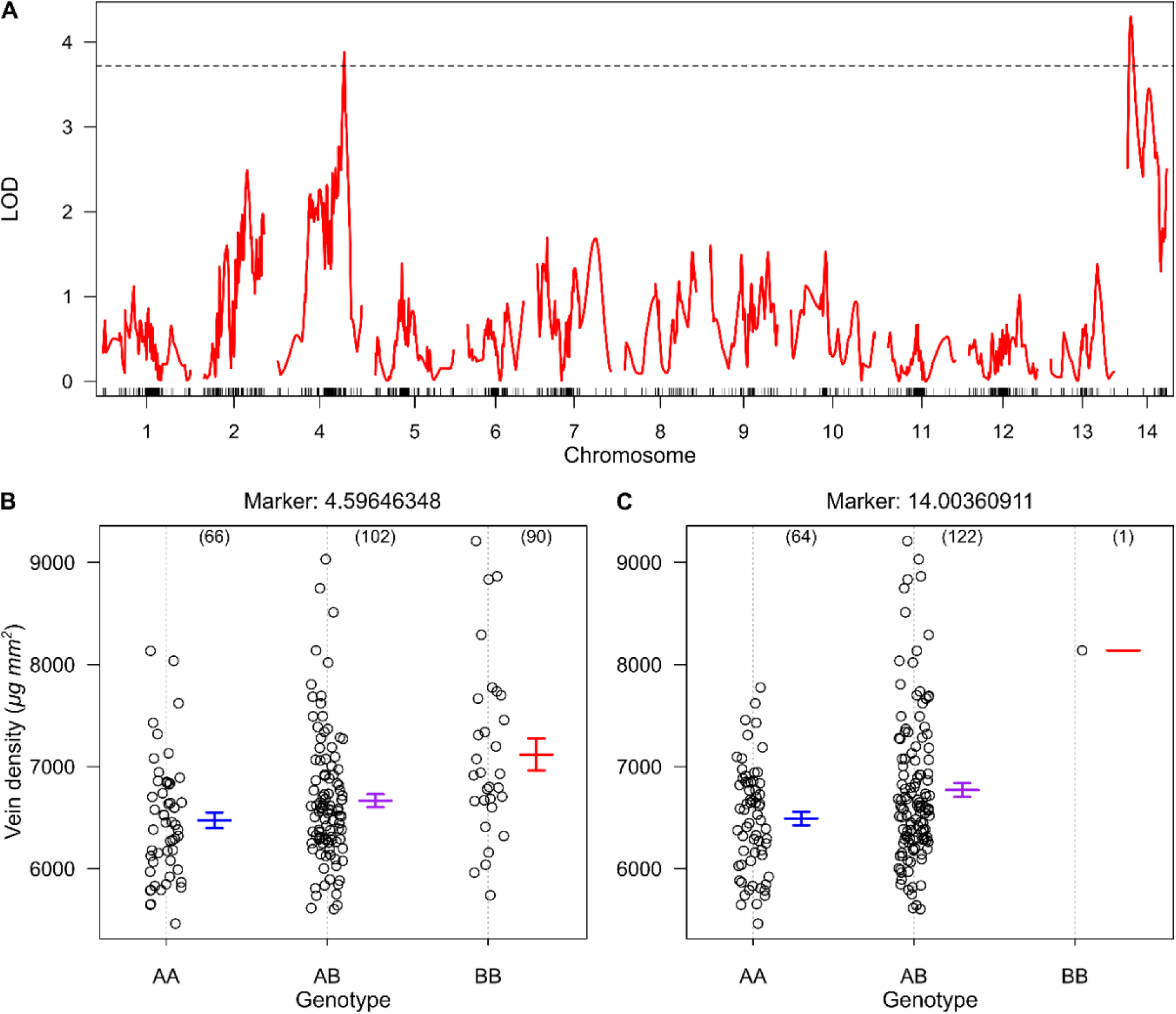
pQTL associated with vein density. **(A)** Single scan of vein density in the Wag19 population with outliers included. **(B)** The genotypic effect of the nearest marker (4.59646348) to the pQTL 4:122.0 and **(C)** the nearest marker (14.00360911) to 14:6.0. Numbers in parentheses are the number of individuals with each genotype. Confidence intervals for average phenotypes within each genotype group are shown as blue for AA, purple for AB, and red for BB.

**Figure S2:**
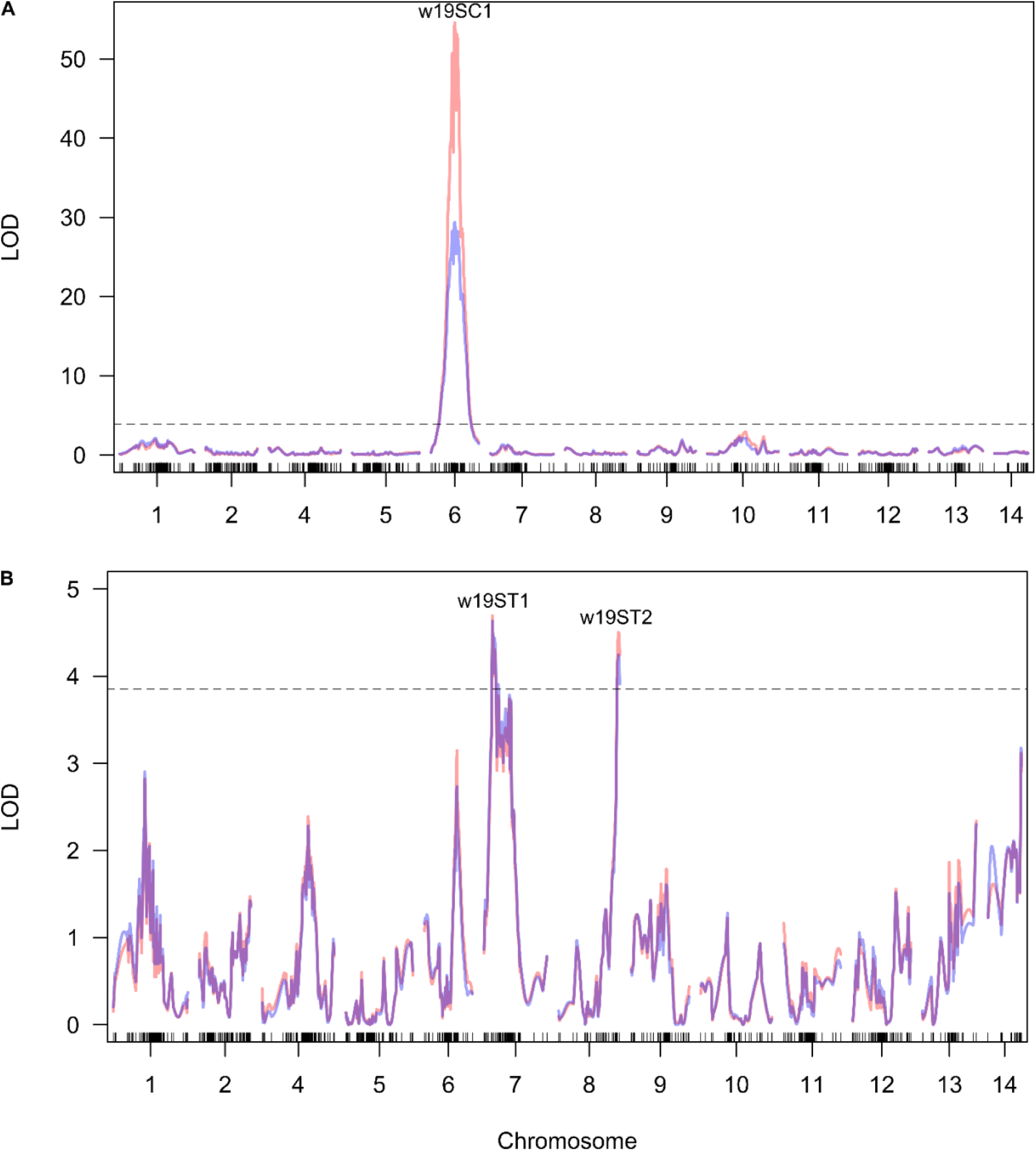
Single QTL scans for (A) stem colour and (B) stem trichome density in Wag19. Red represents the normal model, and blue is the non-parametric model. The dashed line highlights the 0.05 significance threshold after 1000 permutations. The QTL positions are labelled.

**Table S1:** All assessed agronomic traits were significantly different between the parents of the Wag18 population.

|  | Malaysia-03 |  |  | Malawi-02 |  |  |  |  |  |
| --- | --- | --- | --- | --- | --- | --- | --- | --- | --- |
| Trait | n | Mean | S.D | n | Mean | SD | Test | Test stat. | P |
| Violaxanthin Content | 6 | 4.94 | 1.01 | 6 | 12.11 | 2.00 | Parametric | 7.23 | <0.001*** |
| Lutein Content | 6 | 21.00 | 1.36 | 6 | 30.30 | 1.83 | Parametric | 9.36 | <0.001*** |
| Beta-carotene Content | 6 | 50.03 | 3.81 | 6 | 63.56 | 5.89 | Parametric | 4.41 | 0.0017** |
| Total Carotenoid Content | 6 | 75.97 | 4.81 | 6 | 106.00 | 5.97 | Parametric | 9.03 | <0.001*** |
| Alpha-tocopherol Content | 6 | 61.54 | 9.24 | 6 | 35.30 | 9.87 | Parametric | -4.52 | 0.0015** |
| Flowering Time | 6 | 33.33 | 2.16 | 7 | 43.00 | 1.91 | Non-parametric | 42.00 | 0.0025** |
| Plant Height | 6 | 106.35 | 5.60 | 7 | 179.23 | 4.68 | Parametric | 25.61 | <0.001*** |
| Leaf Area | 6 | 40.02 | 7.05 | 7 | 52.38 | 8.09 | Parametric | 2.91 | 0.014* |

n = number of replicates, P = significance value, S.D = Standard Deviation, BSCW = Bundle Sheath Cell Width. Significance is indicated by asterix. Test stat. = Test statistic and is reported as the T or W value for parametric tests and non-parametric tests respectively.

**Table S2:**
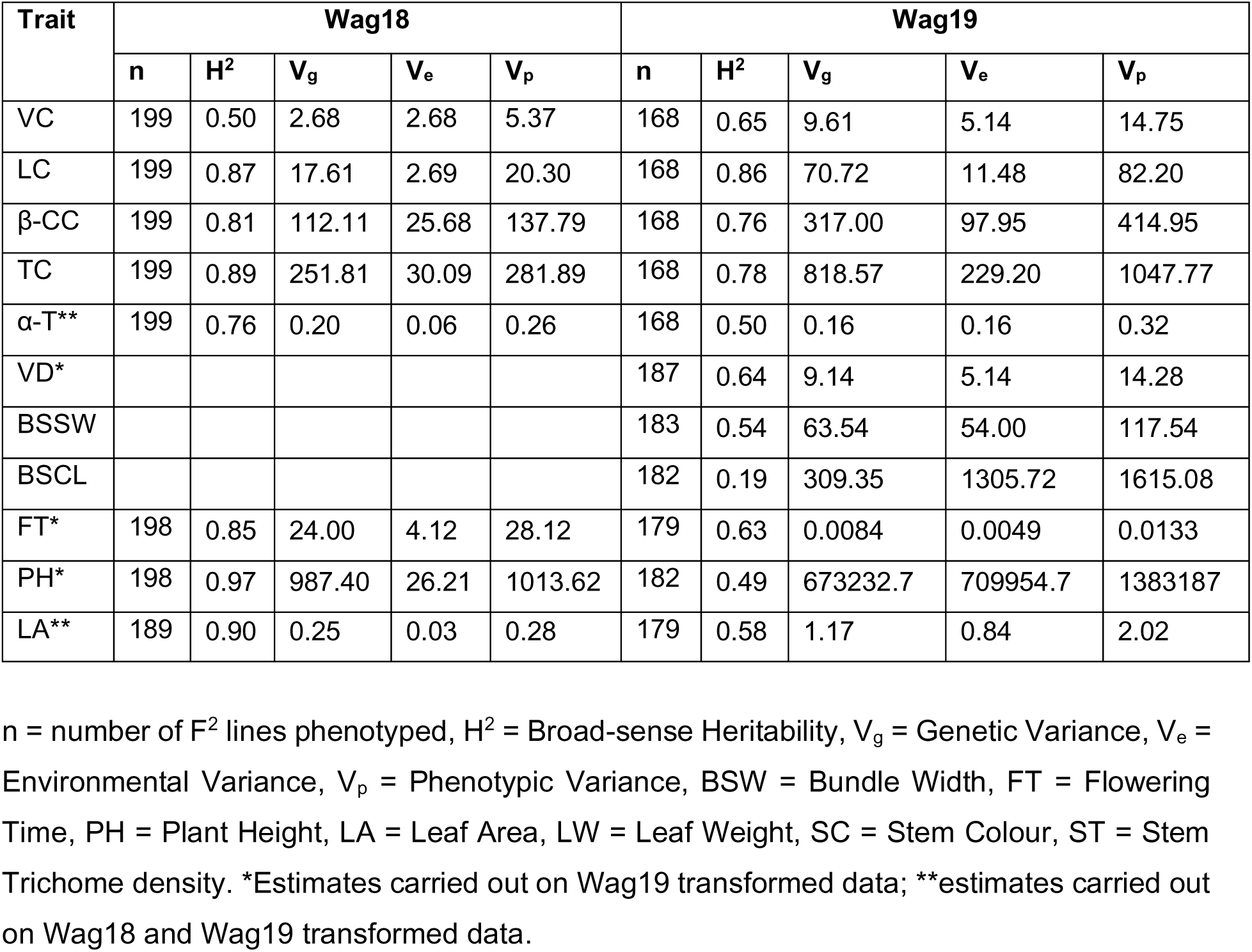
Heritability and variance components for traits in each mapping population.

**Table S3:** Kranz traits, alpha-tocopherol content, flowering time, leaf area, leaf weight, trichome density and stem colour scores were significantly different between the parents while carotene content and plant height were not in the Wag19 population.

|  | Malaysia-01 |  |  | Malawi-01 |  |  |  |  |  |
| --- | --- | --- | --- | --- | --- | --- | --- | --- | --- |
| Trait | n | Mean | S.D | n | Mean | SD | Test | Test stat. | P |
| Violaxanthin Content | 5 | 13.06 | 1.62 | 5 | 16.17 | 2.99 | Parametric | 1.80 | 0.13 |
| Lutein Content | 5 | 31.33 | 3.93 | 5 | 37.15 | 2.34 | Parametric | 2.25 | 0.07 |
| Beta-carotene Content | 5 | 73.56 | 11.37 | 5 | 77.65 | 7.13 | Parametric | 0.54 | 0.61 |
| Total Carotenoid Content | 5 | 117.94 | 16.69 | 5 | 130.97 | 12.45 | Parametric | 1.28 | 0.31 |
| Alpha-tocopherol Content | 5 | 21.71 | 8.24 | 5 | 8.63 | 2.82 | Parametric | -2.58 | 0.049* |
| Vein Density | 5 | 5713 | 267 | 5 | 9822 | 536 | Parametric | 15.35 | <0.001*** |
| BSSW | 4 | 108.76 | 8.48 | 5 | 88.71 | 6.37 | Parametric | -4.07 | 0.0048** |
| BSCL | 4 | 44.51 | 6.55 | 5 | 33.31 | 5.60 | Parametric | -2.77 | 0.028* |
| Flowering Time | 5 | 38.20 | 2.68 | 5 | 52.20 | 2.39 | Non-parametric | 25 | 0.011* |
| Plant Height | 5 | 39.60 | 3.65 | 5 | 45.60 | 13.01 | Parametric | 0.99 | 0.37 |
| Leaf Area | 5 | 18.38 | 1.83 | 5 | 59.57 | 19.77 | Parametric | 4.64 | 0.0094** |
| Stem Trichome Density | 5 | 0.40 | 0.54 | 5 | 1.8 | 0.44 | Non-parametric | 24 | 0.15* |
| Stem Colour | 5 | 0.20 | 0.45 | 5 | 1.6 | 0.54 | Non-parametric | 24 | 0.15* |

n = number of replicates, P = significance value, S.D = Standard Deviation, BSCW = Bundle Sheath Cell Width. Significance is indicated by asterix. Test stat. = Test statistic and is reported as the T or W value for parametric tests and non-parametric tests respectively

**Table S4:** Overall summary of putative QTL identified in the Wag18 population.

| Phenotype | pQTL number | pQTL name (chr:cM) | Nearest marker (chr.bp) | Additive effect | Dominance effect | LOD | Model formula | Model LOD | PVE by model (%) |
| --- | --- | --- | --- | --- | --- | --- | --- | --- | --- |
| Violaxanthin content | Q1 | 1:84.7 | 1.05951559 | -2.78 | 0.83 | 4.98 | y ~ Q1 + Q2 | 7.57 | 16.07 |
| Violaxanthin content | Q2 | 1:89.8 | 1.13366757 | 2.89 | 0.05 | 6.28 | y ~ Q1 + Q2 | 7.57 | 16.07 |
| Lutein content | Q1 | 2:18.0 | 2.2745371 | -2.08 | 0.01 | 4.64 | y ~ Q1 | 4.64 | 10.23 |
| Alpha-tocopherol content* | Q1 | 9:5.7 | 9.00780456 | -0.42 | 0.10 | 17.41 | y ~ Q1 | 17.41 | 33.16 |
| Flowering time | Q1 | 6:45.0 | 6.36025417 | 3.00 | -2.02 | 9.19 | y ~ Q1 | 9.19 | 19.24 |
| Plant height | Q1 | 1:90.8 | 1.13687113 | 25.12 | -1.30 | 12.38 | y ~ Q1 | 12.38 | 25.02 |
| Leaf area* | Q1 | 1:88.9 | 1.11928029 | 0.38 | -0.11 | 8.33 | y ~ Q1 | 8.33 | 18.38 |

*PVE = Percentage of Variance Explained, pQTL name is based on linkage map position: “pChromosome:position(cM)”, nearest marker name is based on physical position: “Chromosome.position(bp)”. Negative and positive additive effects mean the allele from Malaysia -03 and Malawi-02 respectively is responsible for an increase in the trait. \*Analysis carried out on log transformed data.*

**Table S5:** Overall summary of putative QTL identified in the Wag19 population.

| Phenotype | pQTL number | pQTL name (chr:cM) | Nearest marker (chr.bp) | Additive effect | Dominance effect | LOD | Model formula | Model LOD | PVE by model (%) |
| --- | --- | --- | --- | --- | --- | --- | --- | --- | --- |
| Violaxanthin Content | Q1 | 10:58.4 | 10.03027582 | 1.07 | -2.20 | 4.77 | y ~ Q1 | 4.77 | 12.26 |
| Alpha-tocopherol content* | Q1 | 11:122.3 | 11.41198362 | -0.23 | -0.20 | 5.56 | y ~ Q1 | 5.56 | 14.14 |
| Vein Density**** | Q1 | 4:122.0 | 4.59646348 | 1.50 | -0.75 | 3.88 | y ~ Q1 + Q2 | 5.15 | 12.26 |
| Vein Density**** | Q2 | 14:6.0 | 14.00360911 | 4.84 | -3.70 | 4.30 | y ~ Q1 + Q2 | 5.15 | 12.26 |
| Flowering time** | Q1 | 6:77.0 | 6.46920704 | 0.083 | -0.033 | 10.85 | y ~ Q1 | 10.85 | 24.35 |
| Plant height*** | Q1 | 1:83.5 | 1.17961377 | 514.58 | 419.87 | 7.46 | y ~ Q1 + Q2 | 11.62 | 25.47 |
| Plant height*** | Q2 | 2:84.0 | 2.38898522 | -528.14 | 12.25 | 4.48 | y ~ Q1 + Q2 | 11.62 | 25.47 |
| Leaf area**** | Q2 | 1:145.0 | 1.70602349 | -0.27 | 0.99 | 4.98 | y ~ Q1 + Q2 + Q3 | 14.51 | 31.15 |
| Leaf area**** | Q1 | 1:84.3 | 1.18700207 | 0.53 | 0.28 | 4.59 | y ~ Q1 + Q2 + Q3 | 14.51 | 31.15 |
| Leaf area**** | Q3 | 12:73.0 | 12.38068542 | 0.73 | -0.19 | 5.75 | y ~ Q1 + Q2 + Q3 | 14.51 | 31.15 |
| Stem trichomes | Q1 | 7:17.9 | 7.00721601 | -0.26 | -0.08 | 5.18 | y ~ Q1 + Q2 | 9.67 | 21.71 |
| Stem trichomes | Q2 | 8:129.0 | 8.44525256 | 0.25 | 0.15 | 4.98 | y ~ Q1 + Q2 | 9.67 | 21.71 |
| Stem colour | Q1 | 6:51.0 | 6.30828142 | 0.77 | 0.81 | 62.67 | y ~ Q1 + Q2 + Q1:Q2 | 62.75 | 79.56 |
| Stem colour | Q2 | 7:3.9 | 7.00205393 | -0.076 | -0.037 | 8.17 | y ~ Q1 + Q2 + Q1:Q2 | 62.75 | 79.56 |
| Stem colour | Q1:Q2 | NA |  | NA | NA | 6.93 | y ~ Q1 + Q2 + Q1:Q2 | 62.75 | 79.56 |

PVE = Percentage of Variance Explained, pQTL name is based on linkage map position: “pChromosome:position(cM)”, nearest marker name is based on physical position: “Chromosome.position(bp)”. Negative and positive additive effects mean the allele from Malaysia-01 and Malawi-
01 respectively is responsible for an increase in the trait. Analysis carried out on \*log, \*\*cube-root, \*\*\*squared, \*\*\*\*square-root transformed data.

**Table S6:** Summary of ordinal regression on ordinal traits in the Wag19 poulation.

| Trait | pQ1 | pQ2 | Model | AIC | pQ1 L<br>t.value | pQ1 Qu<br>t.value | pQ1 L<br>p.value | pQ1 Qu<br>p.value | pQ2 L<br>t.value | pQ2 Qu<br>t.value | pQ2 L<br>p.value | pQ2 Qu<br>p.value |
| --- | --- | --- | --- | --- | --- | --- | --- | --- | --- | --- | --- | --- |
| Stem<br>trichomes | 7:17.9 | 8:129.0 | y ~ Q1 | 305.28 | -4.25 | 0.34 | <0.001 | 0.73 |  |  |  |  |
| Stem<br>trichomes | 7:17.9 | 8:129.0 | y ~ Q2 | 299.61 |  |  |  |  | 4.03 | -1.02 | <0.001 | 0.31 |
| Stem<br>trichomes | 7:17.9 | 8:129.0 | y ~ Q1<br>+ Q2 | 274.65 | -4.42 | 0.76 | <0.001 | 0.45 | 3.99 | -1.73 | <0.001 | 0.085 |
| Stem<br>colour | 6:51.0 | 7:3.9 | y ~ Q1 | 161.57 | 5.91 | -5.24 | <0.001 | <0.001 |  |  |  |  |
| Stem<br>colour | 6:51.0 | 7:3.9 | y ~ Q2 | 310.21 |  |  |  |  | 0.26 | 0.23 | 0.80 | 0.82 |
| Stem<br>colour | 6:51.0 | 7:3.9 | y ~ Q1<br>+ Q2 | 160.91 | 6.09 | -5.32 | <0.001 | <0.001 | -1.82 | 0.87 | 0.070 | 0.39 |

*pQ = pQTL; L = Linear; Qu = Quadratic. pQTL name is based on linkage map position: "pChromosome:position(cM)". Negative and positive Linear t.values mean the allele from Malaysia-01 and Malawi-01 respectively is responsible for an increase in the trait. Negative and positive Quadratic t.values indicate dominant effects.*

